# Endomicroscopic fluorescence lifetime imaging enables molecular detection and targeted sampling in the distal human lung

**DOI:** 10.64898/2026.08.04.742476

**Authors:** Stuart R Dickson, Erin E Gaughan, Antonella Pellicoro, Bethany Mills, Tarek Haloubi, Mehmet Demirel, Renske Hoekstra, Hazel L Stewart, Lewis Bain, Gareth OS Williams, Adam DL Marshall, Harry AC Wood, Vikki Young, Annya M Bruce, Jean Antonelli, James M Stone, Ahsan R Akram, Tom M Quinn, Thomas Craven, Christopher Haslett, Keith Finlayson, Richard A O’Connor, Manu Shankar-Hari, Kevin Dhaliwal

## Abstract

**Purpose:** Accurate molecular characterisation of infection and inflammation within the distal human lung remains challenging, particularly in critically ill patients, due to limited access to the alveolar space and delayed diagnostic workflows. Molecular imaging approaches capable of real-time detection and targeted sampling could substantially improve diagnostic precision and the future translational development of molecular imaging probes and therapeutics.

**Methods:** In a preclinical setting, we evaluated a clinic-ready endomicroscopic fluorescence lifetime imaging microscopy (eFLIM) platform combined with molecularly targeted SmartProbes for in situ detection of bacteria and activated neutrophils in the distal human lung. A multifunctional 1.9-mm diameter imaging and sampling catheter (Eyes on Target; EoT) enabled real-time fluorescence intensity and lifetime imaging alongside directed alveolar microlavage via a 1.2-mm working channel. Fluorescence intensity and lifetime signatures of Gram-negative bacteria, Gram-positive bacteria, and activated neutrophils were characterised using three wash-free SmartProbes: NBD-PMX, Merocy-Van, and a neutrophil activation probe (NAP). Imaging and sampling performance were assessed in ventilated ex vivo human lungs.

**Results:** EoT reliably navigated to alveolar regions across all lung lobes in both phantom and ventilated human lung models. eFLIM distinguished alveolar microanatomy and enabled probe-specific molecular detection within the distal lung. Increased NBD-PMX signal was detected in *Escherichia coli*–instilled lobes, while Merocy-Van lifetime signatures selectively identified *Staphylococcus aureus*–instilled regions. Activated neutrophils were detected throughout lung tissue following NAP administration. Directed alveolar microlavage enabled recovery of cellular material and bacterial DNA from imaged regions for downstream analysis.

**Conclusion:** eFLIM using EoT combined with molecular SmartProbes enables real-time molecular imaging and targeted sampling within the distal human lung. This platform provides a translatable approach for evaluating infection and inflammation at the alveolar level and supports the clinical development of molecular imaging probes for pulmonary disease.

## Introduction

Pulmonary infection and inflammation are major contributors to morbidity and mortality in critically ill patients, particularly those requiring mechanical ventilation. Pneumonia remains a leading cause of ICU admission and death, with mortality rates approaching 30% in ventilated patients [1, 2]. Despite advances in imaging and microbiological diagnostics, accurately identifying infection within the distal lung remains challenging. Radiological abnormalities are frequently non-specific, and standard microbiological sampling techniques, including bronchoalveolar lavage (BAL), are limited by delayed results and imperfect sensitivity and specificity [3–6]. These limitations contribute to diagnostic uncertainty, delayed treatment optimisation, and excessive use of broad-spectrum antibiotics, accelerating antimicrobial resistance and associated harms [7].

Molecular imaging offers the potential to overcome these limitations by enabling real-time, spatially resolved characterisation of biological processes in situ. Optical endomicroscopy (OEM) has previously demonstrated feasibility for microscopic imaging of the distal lung at the bedside [6, 8]. Using endogenous autofluorescence, OEM allows visualisation of alveolar microstructure; however, its clinical utility has been constrained by limited molecular specificity and an inability to multiplex biological signals. These limitations restrict its capacity to identify pathogens or immune activation directly within the alveolar space.

We hypothesised that molecular characterisation of the distal lung could be enhanced through the integration of three complementary advances: (i) molecularly targeted fluorescent probes capable of selectively labelling pathogens and immune cells; (ii) fluorescence lifetime imaging microscopy (FLIM) to improve contrast, enable multiplexing, and reduce dependence on fluorescence intensity alone; and (iii) a multifunctional catheter capable of both imaging and directed alveolar sampling under direct visualisation.

To achieve molecular specificity, we employed a panel of wash-free SmartProbes designed to generate signal selectively upon interaction with their biological target. These included NBD-PMX for Gram-negative bacterial detection [6], Merocy-Van for Gram-positive bacterial detection [9], and a neutrophil activation probe (NAP) that reports key features of neutrophil activation [10]. FLIM provides an additional imaging dimension by exploiting differences in fluorescence lifetime, which is sensitive to the local molecular microenvironment and largely independent of fluorophore concentration, excitation intensity, and photobleaching [11]. This enables robust multiplexed imaging in complex biological tissues and aligns with emerging molecular imaging strategies in nuclear and optical medicine.

In this study, we combine a clinic-ready fibre-based eFLIM platform (KronoScan) with a fully biocompatible multifunctional imaging and sampling catheter (Eyes on Target; EoT). Using ventilated ex vivo human lungs, we evaluate the ability of this platform to perform real-time molecular imaging of bacteria and activated neutrophils within the alveolar space, while enabling directed sampling for downstream microbiological and molecular analysis. This work provides preclinical validation of a translational molecular imaging approach for characterising distal lung infection and inflammation.

## Materials and Methods

### SmartProbes

All SmartProbes (NBD-PMX, Merocy-Van, and NAP) were synthesised and manufactured as previously described [6, 9, 10]. Probes were diluted to working concentrations in saline: NBD-PMX (11 µM), Merocy-Van (6 µM), and NAP (4 µM).

### Bacterial Preparation

*Staphylococcus aureus* ATCC 25923 and *Escherichia coli* ATCC 25922 were cultured overnight from single colonies in tryptic soy broth (TSB) and Luria–Bertani Lennox broth (LB), respectively, at 37 °C with shaking. Cultures were diluted 1:100 and grown to mid-log phase. For confocal imaging, cultures were harvested at OD₅₉₅ = 1.0, washed in 0.9% NaCl, and resuspended. For lung instillation experiments, bacteria were prepared at OD₅₉₅ = 2.0, washed, and resuspended in 2 mL saline, corresponding to approximately 1 × 10⁸ CFU/mL.

### Benchtop Confocal Laser Scanning Microscopy

Bacterial culture and neutrophil isolation and activation were performed as previously described [6,9,10]. Bacteria and neutrophils were seeded into confocal imaging chambers (ibidi, 80806), pre-coated with poly-D-lysine and human fibronectin.

SmartProbes (100 µL) were added to chambers according to experimental conditions: NAP (4 µM), NBD-PMX (11 µM), or Merocy-Van (6 µM). Confocal fluorescence intensity and lifetime imaging were performed using a Leica SP8 FALCON system under oil immersion with a 63× objective. NAP and NBD-PMX were excited at 488 nm (emission 500–560 nm), while Merocy-Van was excited at 560 nm (emission 570–660 nm). Hybrid (HyD and HyS) detectors were used, and quantification performed using LAS X software (Leica).

### eFLIM Imaging System and EoT Catheter

The design and configuration of the fluorescence lifetime imaging system have been described previously [12, 13]. Briefly, the clinic-ready eFLIM platform (KronoScan) integrates a time-resolved spectrometer (32 lifetime channels) with an achromatic confocal laser scanning microscope. Excitation was provided by a supercontinuum laser (SuperK EVO, NKT Photonics) filtered to 490 nm and 590 nm (Fig. 1a)

**Figure 1.**
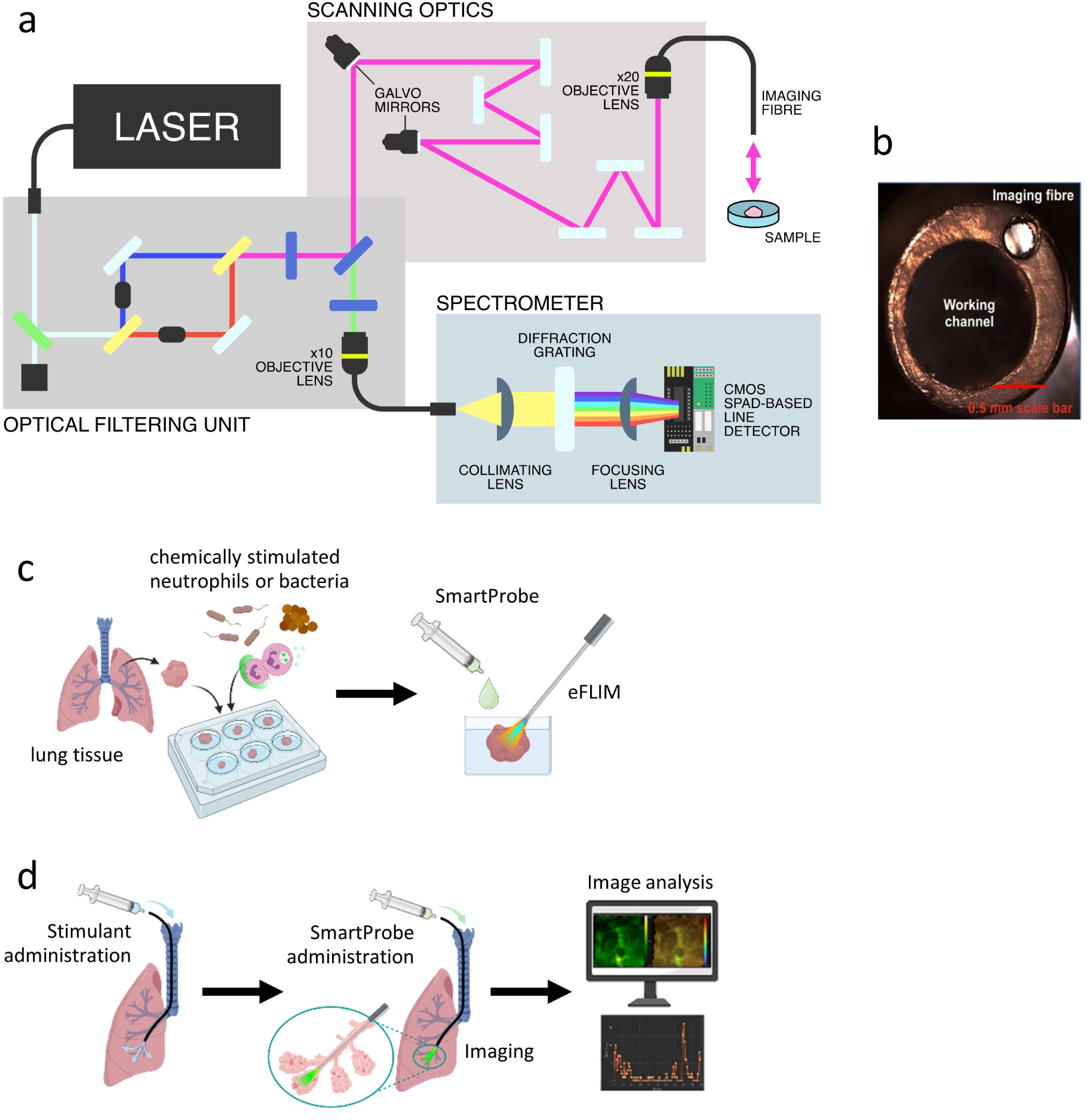
Overview of the eFLIM Imaging System and Experimental Setup. (a) Schematic representation of eFLIM. (b) Image of the end face of the eFLIM fibre Eyes on Target (1.7mm diameter) showing imaging fibre (diameter 350-450 microns) and lumen. Scale bar: 0.5 mm. (c) “Tissue-in-a-Dish” (TiD) setup. 4mm x 4mm x 1mm slices of fresh human lung tissue were placed into individual wells of a black 96-well plate and layered with a cell suspension of either chemically stimulated neutrophils or bacteria. Imaging was then conducted at baseline and after SmartProbe addition. (d) Ex vivo human lung (EVHL) Imaging: Either *E. coli* or *S. aureus* were administered into human lungs, with subsequent eFLIM imaging conducted in treated and untreated lobes post-SmartProbe instillation. c/d: Created in BioRender. Pellicoro, A. (2026) https://BioRender.com/iamojxr

Simultaneous fluorescence intensity and lifetime image sequences (50 frames) were acquired at 128 × 128-pixel resolution (350 × 350 µm field of view), matching the fibre bundle characteristics. Emission spectral bands of 498–565 nm and 620–764 nm were analysed. Lifetime decays were fitted using rapid lifetime determination with a single exponential model, enabling real-time imaging at approximately 3 frames/s with a 20 µs exposure time.

Fully biocompatible multifunctional EoT catheters (outer diameter 1.9 mm, length 2.7m) incorporating an imaging fibre bundle embedded within the catheter wall and a 1.2-mm working channel for fluid delivery and aspiration were used throughout (Fig. 1b). Catheter flexibility and navigability were evaluated in Koken phantom lungs prior to use in human lung studies. The EoT was designed to enable imaging, delivery and aspiration without catheter removal in all regions of the human lung. EoT was also designed as a single use disposable catheter and cost effectiveness enabled by using an embedded imaging bundle designed and developed using low-cost preforms[14]

### Tissue-in-a-Dish Experiments

Human blood was obtained from healthy donors following informed consent. Neutrophils were isolated using Percoll-based density gradients [15] and resuspended at 1 × 10⁷ cells/mL. Cells were activated using calcium ionophore A23187 (1 µM) for 30 minutes before 50 µL aliquots were added to lung tissue wells. Imaging datasets were acquired at baseline and 3 minutes following addition of 50 µL NAP (4 µM in PBS) (Fig. 1c).

For bacterial co-culture experiments, 50 µL of prepared *S. aureus*, *E. coli*, or saline control was added, followed by the appropriate SmartProbe (Merocy-Van or NBD-PMX).

### Ex Vivo Human Lung Tissue

Human lung tissue was obtained from surgical resections under appropriate ethical approval (20/ES/0061 and 15/ES/0094) and stored at −80 °C until use. Tissue was sectioned into approximately 4 × 4 × 1 mm slices and placed into individual wells of black 96-well plates (Thermo Scientific, Microfluor 2) for imaging experiments.

### Ventilated Ex Vivo Human Lung Model

Whole human lungs deemed unsuitable for transplantation were obtained under ethical approval (16/LO/1883 and International Institute for the Advancement of Medicine). Lungs were mechanically ventilated using a Dräger Savina 300 ventilator with tidal volumes of 3–6 mL/kg ideal body weight, fraction of inspired oxygen (FiO₂) of 21%, positive end-expiratory pressure of 5 cmH₂O, pressure support of 5 cmH₂O, and respiratory rate of 12 breaths/min.

Bronchoscopy was performed using a 5.8-mm Ambu® aScope™ 4. Where feasible, lungs were also perfused via pulmonary artery cannulation using a custom ex vivo perfusion circuit. The perfusion system comprised an organ chamber, oxygenator with venous reservoir (Getinge), centrifugal pump (Rotaflow Console, Getinge), and heater (Paratherm). The perfusate consisted of 1 L high-glucose Dulbecco’s Modified Eagle Medium supplemented with 70 g/L bovine serum albumin, 5000 U/L unfractionated heparin, 2.5 g/L Dextran 40, and whole human blood to constitute 4% of perfusate volume.

After a minimum of 2 hours of stabilisation, imaging of the distal lung was performed by advancing the Eyes on Target (EoT) catheter through the bronchoscope working channel. SmartProbes were instilled directly into alveolar regions under visual guidance. Immediately following imaging, 1 mL saline was instilled into the same region and aspirated for downstream microbiological and molecular analysis.

### Simulated Bronchoscopy and Navigation Assessment

To replicate clinical bronchoscopy conditions, the EoT catheter was deployed through the working channel of a commercial bronchoscope (BF-H109, Olympus) within a high-fidelity bronchoscopy training model (LM-092, Koken Co., Japan). Navigation followed a staged approach, beginning in lower lobes and progressing to upper lobes. Once the bronchoscope could advance no further, the EoT catheter was deployed unguided into distal airways.

In ex vivo human lungs, navigation was assessed by advancing the EoT catheter into three subsegments per lobe. Successful transbronchial passage into the alveolar space was confirmed by characteristic alveolar microanatomy visualised on eFLIM imaging (Fig. 2).

**Figure 2.**
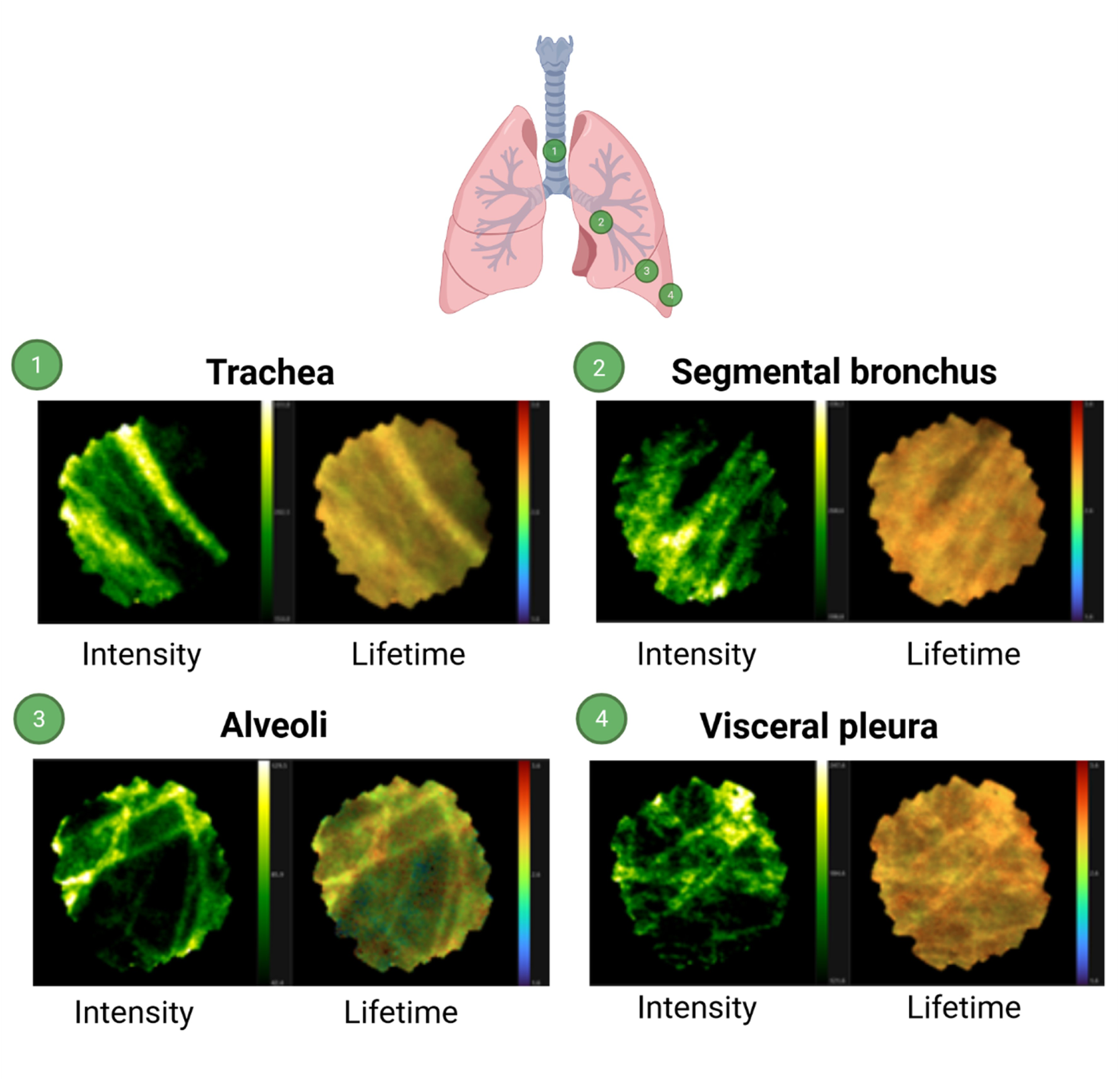
EVHL tissue visualised using eFLIM system. All imaging was obtained with an imaging fibre deployed through the working channel of a bronchoscope in ex-vivo human lungs. (1) Trachea (2) Segmental bronchus (3) Alveoli (4) Visceral pleura. All images were obtained with excitation 490 nm and spectral emission range 498 - 565 nm. For each location, the left-hand image shows fluorescence intensity (autoscaled) and the right-hand image shows fluorescence lifetime (scale 0 - 3.5 ns). Representative image from a single lung. Created in BioRender. Pellicoro, A. (2026) https://BioRender.com/iamojxr

### Ex Vivo Whole Lung Imaging and Sampling

Bacteria were instilled into distinct lung lobes via bronchoscopy using a flexible catheter (1.5 mm APC catheter, Erbe). After 60 minutes, baseline imaging was performed, followed by SmartProbe instillation (NBD-PMX 100 µL, Merocy-Van 200 µL, or NAP 100 µL). Imaging was conducted in 2–3 spatially distinct alveolar regions per lobe. Catheters and bronchosopes were flushed and cleaned with 80% ethanol thoroughly between lobes to prevent cross-contamination (Fig. 1d).

### Alveolar Lavage and Sample Processing

Directed alveolar lavage was performed by instilling up to 2 mL saline via the EoT working channel, followed by a 2 mL air flush and aspiration using a syringe. Lavage samples were cytocentrifuged and stained using Quick-Diff reagents. Slides were scanned and analysed using QuPath software [16]. Remaining lavage samples were processed for CFU enumeration by serial dilution and agar plate colony counting. Additional lavage fluid was processed for bacterial burden by pan-bacterial 16S rRNA real-time PCR, using nucleic-acid extraction from centrifuged lavage supernatant and Ct values as the measure of bacterial load, as previously described [17].

### Image Analysis of data captured by eFLIM in ex vivo lung model

A custom image analysis platform (supplementary methods) incorporating annotation and quantification tools was developed for analysis of NAP, NBD-PMX, and Merocy-Van datasets. Detection algorithms were trained using annotated datasets from independent experiments, with motion-contaminated frames automatically excluded. Algorithmic approaches for probe-specific signal detection are described in supplementary methods.

### Statistical Analysis

Statistical analyses were performed using GraphPad Prism (version 9.5.1). Data are presented as box plots. Comparisons were performed using Mann–Whitney tests, with significance defined as p < 0.05. Associations between culture burden and molecular load were assessed using only aspirates with paired measurements, i.e., samples for which both quantitative culture (CFU/mL) and qPCR (copies/mL) were available. CFU/mL and qPCR copies/mL were log10-transformed and analysed by simple linear regression. The regression equation (slope and intercept), coefficient of determination (R^2^), and the two-sided *p*-value for the null hypothesis that the slope equals zero (F-test) were reported. Samples with zero or non-detect values were excluded from regression analyses due to log transformation.

## Results

### Baseline characterisation of lung anatomy with EoT and eFLIM

An overview of the eFLIM imaging system and experimental set up is shown in Fig. 1. In a high-fidelity lung phantom model, the Eyes on Target (EoT) catheter demonstrated robust flexibility and manoeuvrability, consistently traversing narrow luminal structures representative of distal small airways. Upon exiting simulated distal airways, test samples were introduced to confirm imaging performance under conditions approximating alveolar access (data not shown).

Ex vivo human lung imaging using eFLIM enabled clear delineation of anatomical lung regions based on endogenous autofluorescence. Elastin-rich structures allowed identification of the trachea, main bronchus, segmental bronchi, alveolar spaces, and visceral pleura (Figure 2, Supplementary Fig. S1)). The distinct honeycomb architecture of the alveoli facilitated reliable confirmation of transbronchial catheter passage into the alveolar space, the primary target region for imaging and sampling in suspected pneumonia. Fluorescence lifetime values across these anatomical regions were comparable, ranging from approximately 2.5 to 3.5 ns.

Across three independent experiments using ventilated ex vivo human lungs, EoT consistently achieved transbronchial passage into the alveolar space. Successful alveolar access was confirmed by characteristic alveolar microanatomy visualised on eFLIM imaging. Delivery of 100 µL SmartProbe into the alveolar space was confirmed by transient image distortion and the presence of fluid microbubbles within the imaging field (see supplementary figure S2). These features were observed reproducibly across a minimum of nine lung subsegments from three separate lungs, with a new EoT catheter used in each experiment.

Comparative performance metrics between EoT and the previously described Panoptes fibre [9] are summarised in Table 1, demonstrating improved flexibility, manoeuvrability, and reliability of alveolar sampling with EoT.

Compared with the previously described Panoptes device, EoT demonstrated improved functionality for distal lung interrogation. Panoptes combined an imaging fibre with two small capillary channels for probe delivery, whereas EoT integrates an imaging fibre with a larger 1.2-mm working channel. This configuration enabled not only SmartProbe delivery but also directed alveolar lavage and aspiration from the same region. Across 26 attempted alveolar lavages, EoT enabled fluid recovery in 23 procedures (88%), with aspirated volumes ranging from 115 to 586 µL. These data demonstrate that EoT extends fibre-based molecular alveoscopy from probe delivery and imaging towards combined imaging, delivery, and spatially matched sampling.

### Detection of activated neutrophils using fluorescence lifetime imaging

Activated neutrophils labelled with the neutrophil activation probe (NAP) exhibited characteristic fluorescence intensity and lifetime signatures in vitro. Confocal imaging demonstrated increased punctate fluorescence intensity within activated neutrophils (Figure 3a), consistent with prior reports [10]. In tissue-in-a-dish experiments, NAP-labelled neutrophils demonstrated both increased fluorescence intensity and reduced fluorescence lifetime relative to background lung tissue (Figure 3b / c).

**Figure 3.**
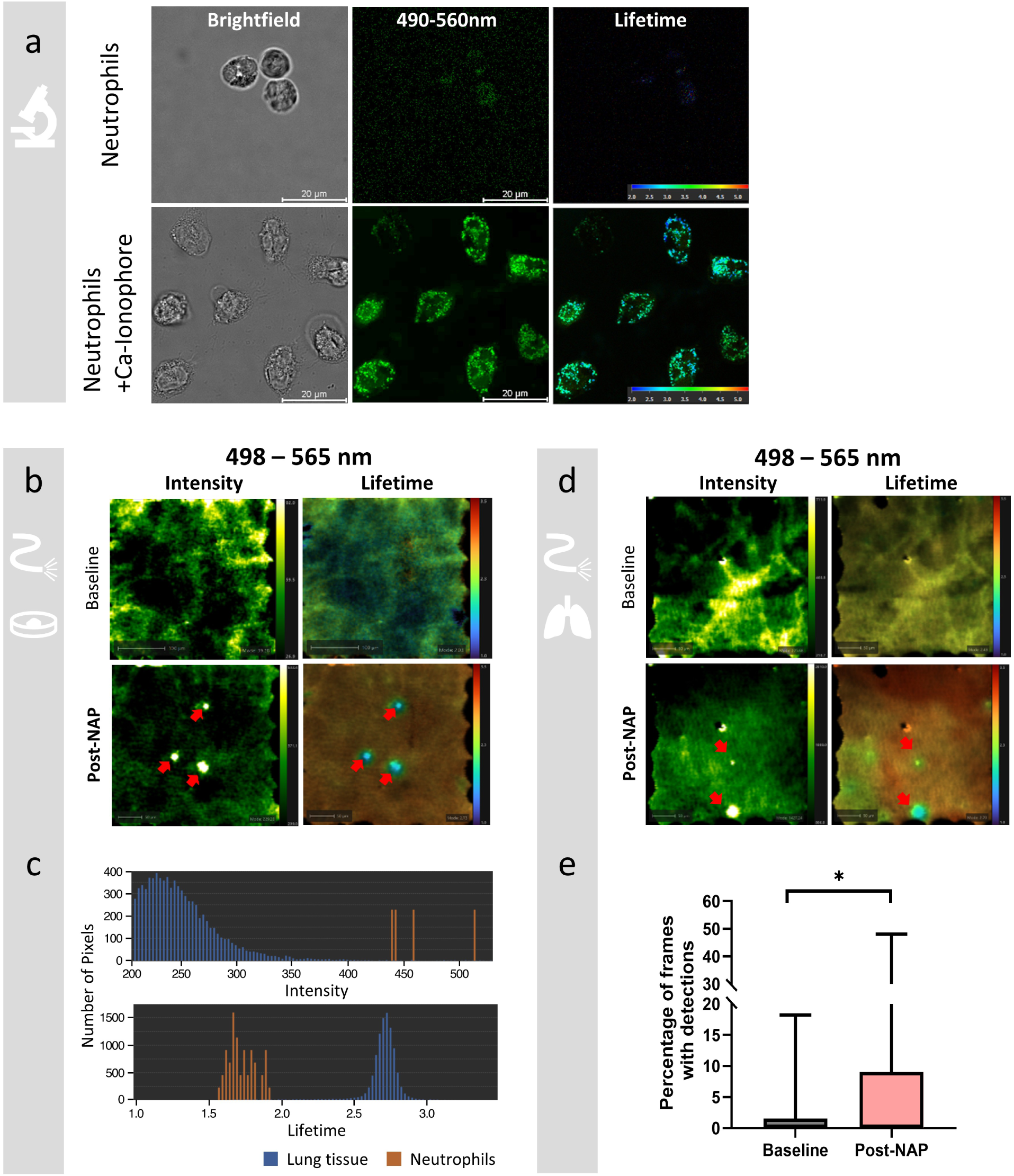
eFLIM system and NAP can identify activated neutrophils within the ex-vivo human lung (EVHL). (a) Representative images of human neutrophils, stimulated with Ca lonophore (A23187) (lµM) and imaged on Stellaris 8 Falcon Flim Microscope (Ex: 488, Em: 490-565 nm. (b) Representative frames of TiD imaging obtained with eFLIM. Activated neutrophils (red arrows) incubated with human lung tissue at baseline and following administration of 50 µL NAP (4 µM). Intensity range for each image is autoscaled by the in-house image analysis platform. Lifetime range is 1.0 - 3.5. (c) Representative histogram plots of neutrophils (orange) and background lung tissue (blue) obtained from the single exemplary imaging frame seen in b (Post-NAP). Neutrophils are indicated by red arrows. (d) Representative frames from EVHL ventilated and perfused lungs pre- (Baseline) and immediately post-delivery of 100 µL (4 µM, Post-NAP). Neutrophils are marked with red arrows in intensity and lifetime images. (e) Analysis of EVHL ventilated and perfused NAP data (2x 100 frames in 3 separate subsections of each lung in 4 independent experiments). Box plot representing the percentage of frames with NAP detections (as per imaging analysis algorithm, based on intensity and LT signatures) in each section imaged. Statistical analysis performed by unpaired students t-test, whiskers extend to the minimum and maximum values, * = p < 0.05.

In ventilated ex vivo human lungs, automated detection algorithms identified activated neutrophils based on regions of elevated intensity and reduced lifetime relative to background autofluorescence (Figure 3d / Supplementary Fig. S3 / S5). Analysis of 100-frame datasets across 12 lung subsegments from four independent experiments demonstrated a significant increase in frames containing NAP-positive signal following probe administration compared with baseline (9.6% vs. 1.0%, p = 0.03; Figure 3e). These findings confirm the feasibility of detecting activated neutrophils within the alveolar space using eFLIM.

### Gram-negative bacterial detection using NBD-PMX

Fluorescence intensity and lifetime confocal images of NBD-PMX labelled Gram-negative *E. coli* and Gram-positive *S. aureus* are shown in Fig. 4a. NBD-PMX has previously enabled *in situ* detection of Gram-negative bacteria using intensity-based optical endomicroscopy [6]. However, spectral overlap with NAP precludes multiplexed intensity-only imaging. Incorporation of fluorescence lifetime imaging enables separation of these signals within a multiplexed framework.

**Figure 4.**
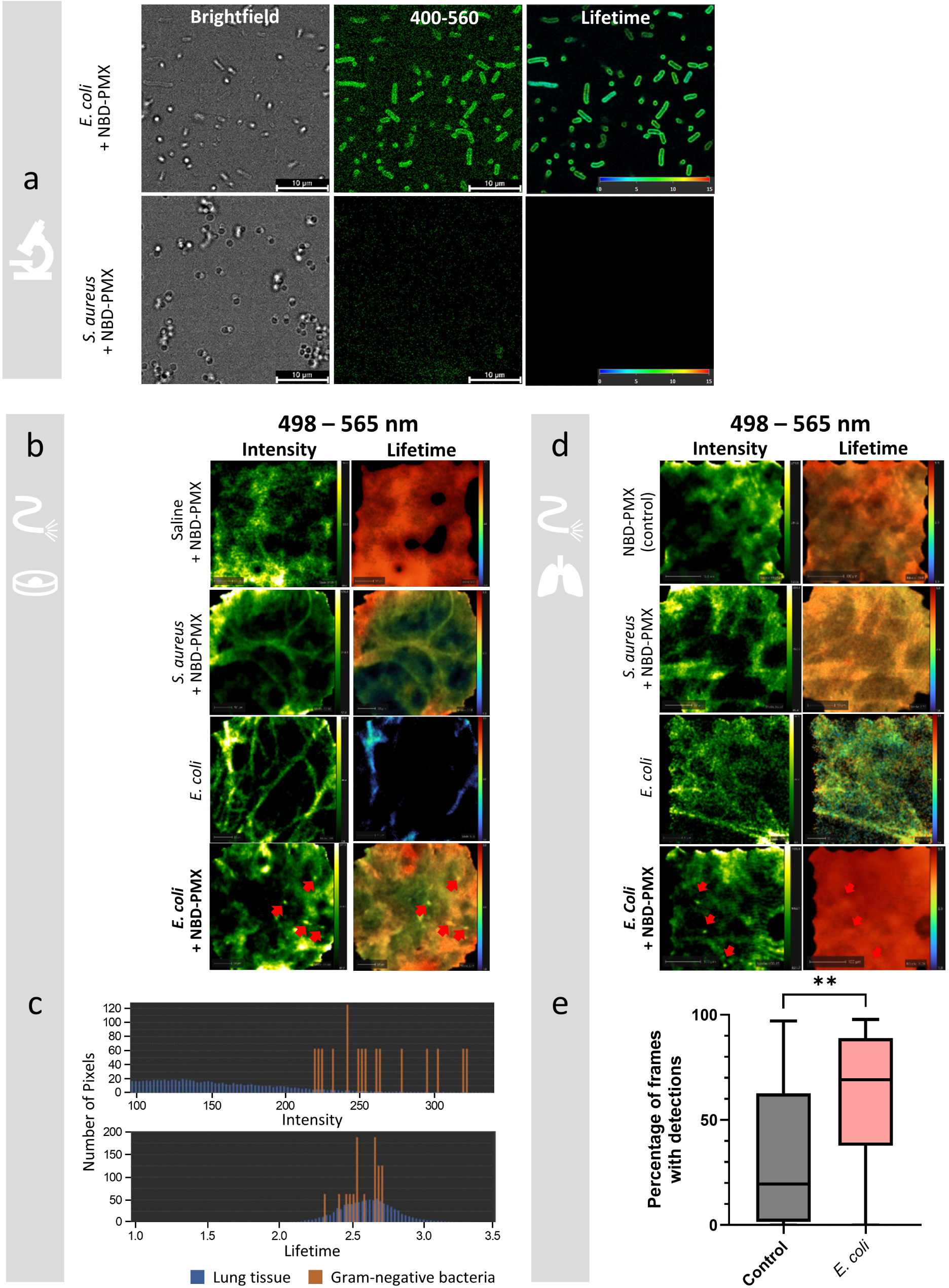
eFLIM system and NBD-PMX can identify gram-negative bacteria within the EVHL. (a) Representative images from a Stellaris 8 Falcon Flim Microscope of *E. coli* and 5. *aureus* incubated with NBD-PMX (Ex: 488, Em: 400-560nm). (b) Representative frames from TiD experiments imaged with eFLIM with target (*E. coli)* and off target (saline, 5. *aureus)* additions pre-and post-NBD-PMX (lluM). Intensity range for each image is autoscaled by the image analysis system. Lifetime range is 1.0 - 3.5. Labelled *E. coli* are identified by red arrows. (c) Representative histogram plots of labelled gram-negative bacteria (orange) and background lung tissue (blue) obtained from single exemplary imaging frame seen in B (*E. coli* + NBD-PMX). (d) Representative eFLIM frames from ventilation and perfusion EVHL experiments. Comparing untreated (control) lobes to target *(E.coli)* and off-target (*S. aureus)* treated lobes with and without NBD-PMX (11 mM). Labelled *E. coli* are marked with red arrows in both fluorescence intensity and lifetime images. (e) Analysis of EVHL NBD-PMX data (2x 100 frames from 3 separate subsections of each lung in 4 independent experiments). Box plot represents the percentage of frames with NBD-PMX detections (as per imaging analysis algorithm, based on intensity signatures) in each section imaged. Statistical analysis performed by unpaired students t-test, whiskers extend to the minimum and maximum values.**= p < 0.01.

In tissue-in-a-dish experiments and ventilated ex vivo human lungs, NBD-PMX-labelled bacteria exhibited a characteristic transient “blinking” behaviour, with brief increases in fluorescence intensity relative to background lung autofluorescence (Figure 4b, c, d and Supplementary Fig. S4). While the fluorescence lifetime of NBD-PMX-labelled bacteria was not distinguishable from background tissue, regions exposed to NBD-PMX exhibited altered lifetime distributions compared with naïve lung tissue, effectively providing a “painting” signature that confirmed probe delivery.

Analysis using a bespoke detection algorithm demonstrated a significantly higher proportion of frames containing blinking signatures in *E. coli*–instilled lobes compared with control lobes across four independent experiments (61.7% vs. 33.3%, p = 0.001; Figure 4e). These data demonstrate that eFLIM enables detection of Gram-negative bacterial signatures within SmartProbe-exposed regions of the distal lung.

### Gram-positive bacterial detection using Merocy-Van fluorescence lifetime signatures

Merocy-Van selectively labels Gram-positive bacteria by incorporation into the bacterial cell wall [9]. Fluorescence intensity and lifetime confocal imaging of Merocy-Van labelled *S. aureus* and *E. coli* is shown in Fig. 5a. Following Merocy-Van administration in both tissue-in-a-dish and ex vivo lung models, comparable fluorescence intensity was observed in both *S. aureus*–instilled and control regions, indicating non-specific tissue fluorescence in the intensity channel and providing a “painting” signature. In contrast, fluorescence lifetime analysis revealed marked differences between infected and control regions (Fig. 5b).

**Figure 5.**
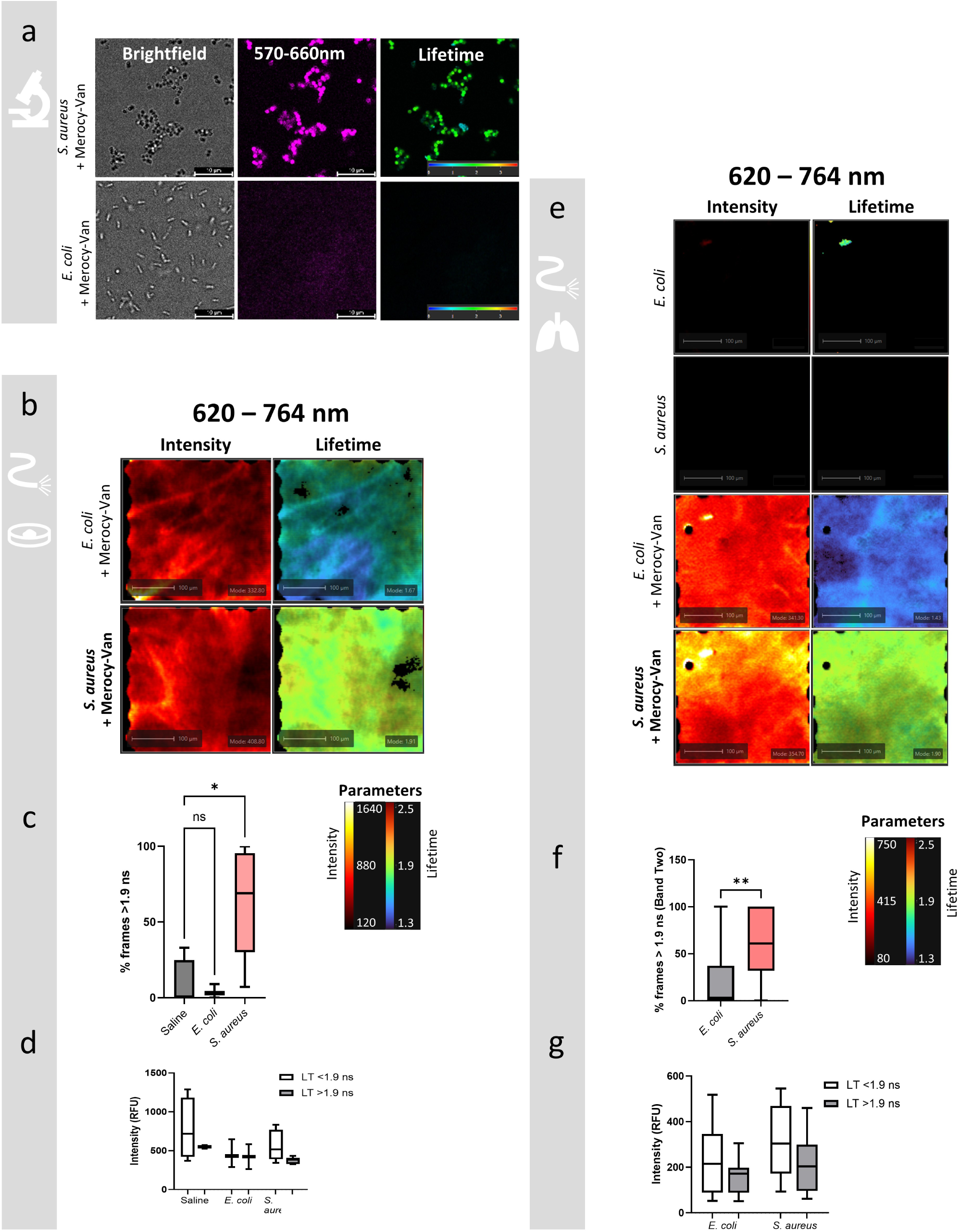
eFLIM system and Merocy-Van can identify Gram-positive bacteria within the EVHL. (a) Representative images of *S. aureus* and *E.coli* incubated with Merocy-Van (Ex:590, Em: 570-660 nm) using a Stellaris 8 Falcon Flim Microscope. (b) Representative imaging frames from TiD experiments with target (*S. aureus*) and off-target (*E. coli*) bacteria with Merocy-Van (6 µM). Data collected via eFLIM with Merocy-Van signatures shown in 620-764 nm lifetime. All displayed data is scaled consistently across each condition per imaging parameter. (c) % of imaged frames with fluorescence LT (620 - 764 nm) above 1.9 ns from tissue assays. (d) Quantification of fluorescence intensity (620 - 764 nm) from tissue assays. EVLH tissue data presented from 4 independent experiments. (e) Representative imaging frames from ventilated EVHL experiments with target (*S. aureus*) and off-target (*E. coli*) bacteria with (and without) Merocy-Van (6 µM). Data collected via eFLIM, with Merocy-Van signatures shown in 620 - 764 nm lifetime. All displayed data is scaled consistently across each condition per imaging parameter. F) % of imaged frames with 620-764 nm fluorescence lifetime above 1.9 ns from ventilated EVHL assays. G) Quantification of 620 - 764 nm fluorescence intensity from the ventilated EVHL assays. Data presented from 4 independent experiments, *S. aureus* and *E. coli* data from 20 imaging videos for each condition. Each video is represented by a single point within the box plot. TiD statistical analysis performed by one-way ANOVA, EVHL statistical analysis performed by unpaired students t-test. Whiskers extend to the minimum and maximum values. *P<0.05; **P<0.001.

In both experimental models, *S. aureus*–instilled regions exhibited significantly altered fluorescence lifetime distributions compared with controls (Figure 5b-f). The observed lifetime values in tissue were shorter than those measured in vitro (approximately 2.0 ns vs. 3.25 ns), consistent with microenvironmental effects on fluorophore lifetime. Based on these observations, a lifetime threshold of ≥1.9 ns was selected to classify Merocy-Van– positive frames. Using this threshold, approximately 60% of *S. aureus*–predicted frames were classified as positive in both tissue-in-a-dish and ex vivo lung experiments, compared with fewer than 10% of negative frames in tissue-in-a-dish experiments (p = 0.0213) and fewer than 20% in ex vivo lungs (p = 0.0019) (Figure 5d, g). Importantly, fluorescence lifetime signatures were independent of fluorescence intensity (Figure 5c, f), supporting robust molecular discrimination of Gram-positive bacterial infection.

### EoT enables bacterial aspiration and PCR of alveolar samples

EoT sampling enabled collection of alveolar aspirates suitable for downstream bacterial quantification by qPCR (Fig. 6). Among 109 aspirates tested per target, qPCR detected bacterial DNA across a broad dynamic range (typically 10^4^ to ≥10^7^ copies/mL; Fig. 6a and c). S. aureus was most frequently detected (80/109, 73.4%), followed by E. coli (58/109, 53.2%); Detection was dominated by *S. aureus* (80/109, 73.4%) and *E. coli* (58/109, 53.2%), reflecting the inclusion of *S. aureus*-/ *E. coli*-instilled samples in the dataset. Less frequent detections were observed for A. baumannii (21/109, 19.3%), Enterobacter cloacae complex (17/109, 15.6%), P. aeruginosa (17/109, 15.6%), H. influenzae (16/109, 14.7%), S. agalactiae (15/109, 13.8%), K. pneumoniae (11/109, 10.1%), and S. pneumoniae (2/109, 1.8%).

**Figure 6.**
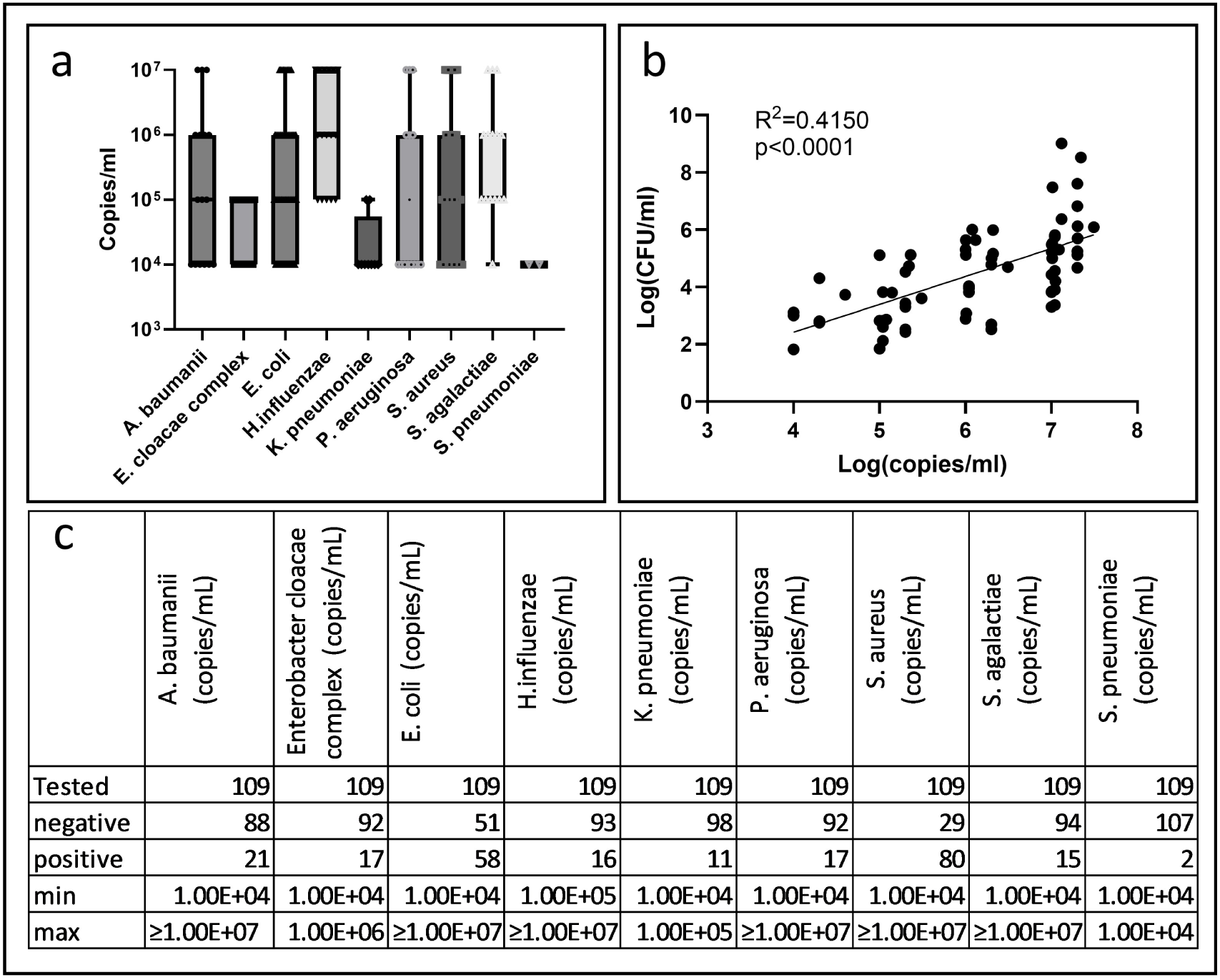
EoT enables bacterial aspiration and PCR of alveolar samples. A) Box plots of qPCR copy number per ml for each bacterial species detected in EoT-collected alveolar aspirates. Boxes indicate the median and interquartile range, whiskers show the spread of the data, and overlaid points represent individual samples. Additionally, the following microorganisms were tested but always found below level of detection (<l0^−4^): *K. aerogenes; K. oxytoca; M. catarrhalis; Proteus spp.;* *S. marcescens;* *S. pyogenes;* Adenovirus, Influenza B virus; Metapneumovirus; Parainfluenza virus; Respiratory Syncytial Virus; Rhinovirus/Enterovirus; Seasonal Coronavirus; MERS-CoV; *Chlamydia pneumoniae; Legionella pneumophi/a; Mycop/asma pneumoniae*. B) Correlation between culture and molecular quantification in matched aspirates shown on log10 scales. Linear regression on log10-transformed CFU/ml and qPCR copies/ml; R^2^ and the two-sided *p*-value for the null hypothesis that the slope equals zero are shown. C) Table summarising the dataset: total number of aspirates analysed, numbers of qPCR-positive and qPCR-negative samples, and (for qPCR-positive samples) the minimum and maximum copy numbers per ml.

Minimum detected loads were 10^4^ copies/mL for all targets except H. influenzae (10^5^ copies/mL), while maximum loads reached ≥10^7^ copies/mL for A. baumannii, E. coli, H. influenzae, P. aeruginosa, S. aureus, and S. agalactiae (lower maxima: Enterobacter cloacae complex 10^6^ copies/mL; K. pneumoniae 10^5^ copies/mL; S. pneumoniae 10^4^ copies/mL). Additional targets—K. aerogenes, K. oxytoca, M. catarrhalis, Proteus spp., S. marcescens, S. pyogenes, adenovirus, influenza B virus, metapneumovirus, parainfluenza virus, respiratory syncytial virus, rhinovirus/enterovirus, seasonal coronavirus, MERS-CoV, Chlamydia pneumoniae, Legionella pneumophila, and Mycoplasma pneumoniae—were tested but remained below the assay limit of detection in all aspirates (all < 10^4^ copies/mL). In matched samples with quantifiable results for both modalities, culture burden correlated positively with qPCR load on log10-transformed scales (linear regression: Y = 0.9749X − 1.483; R^2^ = 0.415; p < 0.0001; Fig. 6b).

Together, these results demonstrate that eFLIM combined with molecular SmartProbes enables real-time molecular imaging of activated neutrophils, Gram-negative bacteria, and Gram-positive bacteria within the distal human lung, while supporting targeted alveolar sampling from the same anatomical region

## Discussion

In this study, we demonstrate the feasibility of real-time molecular imaging and targeted sampling within the distal human lung using a clinic-ready endomicroscopic fluorescence lifetime imaging microscopy (eFLIM) platform combined with molecularly targeted SmartProbes and a multifunctional imaging and sampling catheter. This approach enables in situ detection of bacteria and activated neutrophils at the alveolar level, while simultaneously supporting directed alveolar sampling for downstream microbiological and molecular analysis. Together, these capabilities address a major unmet need in pulmonary molecular imaging: the ability to characterise infection and inflammation directly within the distal lung in real time.

The integration of fluorescence lifetime imaging with molecularly targeted probes represents a key advance over prior intensity-based optical endomicroscopy approaches. Fluorescence lifetime imaging provides contrast that is largely independent of fluorophore concentration, excitation intensity, and photobleaching, while remaining sensitive to the local molecular microenvironment [18, 19]. In the context of pulmonary imaging, where tissue heterogeneity, motion, and variable probe distribution pose substantial challenges, lifetime-based contrast enables more robust signal discrimination and multiplexing. By combining spectral and fluorescence lifetime information, eFLIM provided multidimensional contrast for SmartProbe detection. NAP and NBD-PMX signals were acquired within the 498–565 nm emission band but were differentiated by their fluorescence lifetime, morphology, and temporal dynamics, whereas Merocy-Van was detected in the 620–764 nm emission band and identified primarily through its lifetime signature. This enabled the same imaging platform to distinguish activated neutrophils, Gram-negative bacteria, and Gram-positive bacteria in distal human lung tissue.

The Eyes on Target (EoT) catheter further extends the translational relevance of this platform by enabling combined imaging and sampling and delivery through a single, fully biocompatible device. Unlike prior fibre-based approaches that required separate catheters for imaging and lavage [6], EoT allows probe delivery, real-time visualisation, and aspiration from the same alveolar region. This design minimises the risk of sampling from anatomically distinct regions, reduces procedural complexity, and mitigates contamination from proximal airways. The ability to reliably perform directed alveolar lavage under direct visual guidance represents a substantial advance for correlating molecular imaging signals with microbiological and cellular analyses.

The key technical advance of EoT over Panoptes is the integration of a clinically useful working channel for directed sampling. Panoptes was designed primarily for optical imaging and local SmartProbe delivery, using two small capillary channels with an internal diameter of 326 µm. Although these capillaries permitted fluid delivery and limited sample removal, they were not suited to reliable alveolar microlavage. EoT addresses this limitation by incorporating a 1.2-mm working channel, enabling both probe delivery and aspiration from the same imaged alveolar region.

The detection of activated neutrophils using the neutrophil activation probe (NAP) highlights the ability of eFLIM to interrogate immune activity within the distal lung. Activated neutrophils are central to the pathogenesis of pneumonia, acute respiratory distress syndrome, and other inflammatory lung diseases. The distinct fluorescence lifetime and intensity signatures observed in both tissue-in-a-dish and ex vivo lung models demonstrate that immune activation can be detected directly within the alveolar space. Importantly, the presence of NAP signal in control lobes underscores the complex baseline inflammatory milieu of the human lung and aligns with emerging understanding of pulmonary immune surveillance and resident immune populations [20].

Similarly, the detection of bacterial signatures using NBD-PMX and Merocy-Van demonstrates the complementary roles of intensity-based and lifetime-based contrast in molecular imaging. For Gram-negative bacteria, the characteristic transient intensity “blinking” signature of NBD-PMX enabled detection within SmartProbe-exposed regions, supported by lifetime-based confirmation of probe distribution. For Gram-positive bacteria, fluorescence lifetime provided the primary discriminant, with Merocy-Van lifetime shifts enabling selective identification of *Staphylococcus aureus* despite non-specific intensity signals. These findings emphasise the value of fluorescence lifetime imaging for molecular discrimination in complex biological environments and align with broader trends in molecular imaging towards multiparametric signal extraction.

Image analysis remains a critical component of translating eFLIM into robust clinical workflows. In this study, bespoke rule-based algorithms enabled automated detection of probe-specific signatures, but were inherently sensitive to motion artefacts and system noise. Motion artefacts are a well-recognised challenge in pulmonary optical imaging [21], particularly in ventilated or spontaneously breathing lungs. While the current algorithms provide proof-of-concept validation, future work will focus on integrating machine learning approaches to improve robustness, automate artefact rejection, and enhance real-time detection performance. Recent advances in machine learning applied to fluorescence lifetime imaging have demonstrated substantial gains in accuracy and computational efficiency [22], and preliminary work using unsupervised approaches has already shown promise in reducing false-positive detections for bacterial imaging [23].

A technical limitation of the current eFLIM configuration is the use of reduced temporal binning to maintain real-time imaging at clinically relevant frame rates. While the system is capable of acquiring higher temporal resolution lifetime data, increased acquisition times compromise real-time visualisation and anatomical navigation [24]. Optimising the balance between temporal resolution, frame rate, and signal fidelity will be an important focus of future system development, particularly as more complex molecular probe panels are introduced.

Beyond infection imaging, this platform opens new opportunities to study distal lung biology more broadly. The alveolar duct and alveolar space remain underexplored in vivo due to limited accessibility with conventional imaging and sampling tools. The combination of lifetime-resolved confocal imaging with targeted sampling and precision delivery under direct vision provides a unique opportunity to correlate molecular imaging signals with cellular, microbial, and molecular readouts from precisely defined anatomical regions. Such an approach could support investigation of spatial heterogeneity in lung injury, immune responses, and treatment effects, with potential applications extending beyond infection to interstitial lung disease, acute respiratory distress syndrome, and therapy response assessment.

In conclusion, we demonstrate that eFLIM combined with molecular SmartProbes and a multifunctional imaging and sampling catheter enables real-time molecular imaging and targeted sampling within the distal human lung. This work provides preclinical validation of a translational molecular imaging platform capable of detecting bacterial infection and immune activation at the alveolar level. These findings support further clinical evaluation of eFLIM-based approaches for characterising distal lung pathology and for the development and validation of novel molecular imaging probes in pulmonary disease.

## Supplementary Methods

### Rule-based Detection Algorithms for SmartProbe Signal Identification

#### Overview

Custom rule-based image analysis algorithms were developed to enable automated detection of probe-specific fluorescence signatures associated with activated neutrophils (NAP), Gram-negative bacteria (NBD-PMX), and Gram-positive bacteria (Merocy-Van) in endomicroscopic fluorescence lifetime imaging microscopy (eFLIM) datasets. Algorithms were designed to operate on fluorescence intensity and fluorescence lifetime (FLT) data acquired in real time, while minimising false-positive detections arising from motion artefacts, noise, or background autofluorescence.

All algorithms were implemented within an in-house image analysis platform incorporating annotation, visualisation, and quantification tools.

#### Dataset Curation and Training

Algorithm development utilised a training dataset comprising four independent ventilated ex vivo human lung experiments. Frames affected by excessive motion or interference were automatically excluded prior to analysis. Specifically, frames in which more than 30% of pixels exhibited a greater than 30% change in green-channel intensity relative to the preceding frame were excluded.

Manual annotations were independently performed by four researchers. Annotations identified:

- Activated neutrophils for NAP datasets
- Gram-negative bacterial signatures for NBD-PMX datasets

These annotations informed numerical characterisation of fluorescence intensity and lifetime features and guided selection of detection thresholds. Algorithm performance was subsequently evaluated on independent datasets.

#### Activated Neutrophil Detection (NAP)

NAP-labelled activated neutrophils were characterised by regions exhibiting high fluorescence intensity and reduced fluorescence lifetime within the 498–565 nm emission band.

Because NAP-positive regions vary in size and morphology, a clustering-based flood-fill approach was used.

Detection Procedure

For each frame:

1. Each pixel was evaluated sequentially:

- Pixels with fluorescence lifetime exceeding an absolute threshold of 2.5 ns or exceeding the 20th percentile of the frame’s lifetime distribution were excluded.
- Pixels with fluorescence intensity below the 98th percentile of the frame’s intensity distribution were excluded.
2. Pixels satisfying both criteria were assigned to a cluster.
3. Orthogonally adjacent pixels were recursively evaluated using the same criteria and added to the cluster if eligible.
4. Clusters containing fewer than five pixels were excluded to reduce false-positive detections arising from isolated noise pixels.
5. Accepted clusters were classified as NAP-positive regions.

Frames containing one or more NAP-positive clusters were classified as positive detection frames.

#### Gram-negative Bacterial Detection (NBD-PMX)

NBD-PMX-labelled Gram-negative bacteria were identified based on a characteristic transient “blinking” fluorescence intensity signature in the 498–565 nm emission band. These signals typically appeared as single-pixel or small-region events persisting for one to three frames.

Detection Procedure

1. For each pixel coordinate across consecutive frames:

- Pixels exceeding the 99th percentile of fluorescence intensity were excluded to prevent detection of bright artefacts.
2. For remaining pixels:

- The frame-to-frame fluorescence intensity difference (ΔI) was calculated.
- A candidate detection was flagged if:

- ÄI exceeded +42.5 units between two consecutive frames, and
- ÄI subsequently decreased by −42.5 units within one frame.
3. Only pixel coordinates satisfying both conditions were classified as NBD-PMX-positive detections.

This bidirectional thresholding approach was designed to reduce false-positive detections caused by noise or gradual intensity fluctuations.

Frames containing one or more blinking events were classified as positive detection frames.

#### Gram-positive Bacterial Detection (Merocy-Van)

Merocy-Van detection relied primarily on fluorescence lifetime contrast rather than intensity, due to non-specific tissue fluorescence observed in the intensity channel.

Pre-processing

- Images were processed using fixed brightness and contrast settings optimised for intensity visualisation.
- Fluorescence lifetime maps were displayed using a fixed lifetime range.

#### Frame Inclusion Criteria

Frames were excluded from analysis if the mean fluorescence intensity within the Merocy-Van spectral band was below:

- 50 relative fluorescence units (RFU) for tissue-in-a-dish experiments
- 140 RFU for ventilated ex vivo lung experiments

These thresholds were used to exclude frames lacking tissue contact. Detection Procedure

1. For included frames, a fluorescence lifetime threshold of ≥1.9 ns was applied within the Merocy-Van spectral region.
2. Frames in which the average lifetime exceeded this threshold were classified as Merocy-Van-positive.
3. For each experimental repeat, the percentage of positive frames was calculated and compared between *Staphylococcus aureus*–instilled and control conditions.

Lifetime-based classification was independent of fluorescence intensity, enabling discrimination of Gram-positive bacterial signatures in the presence of background tissue fluorescence.

#### Statistical Considerations

Detection outcomes were quantified as the proportion of frames classified as positive within each dataset. Statistical comparisons between experimental conditions were performed as described in the main Methods section.

## Supporting information

FigS1_Static with description

FigS2_Static with description

FigS5_Static with description

## References

1. Ferreira-Coimbra, J., C. Sarda, and J. Rello, Burden of Community-Acquired Pneumonia and Unmet Clinical Needs. Advances in Therapy, 2020. 37(4): p. 1302–1318.

2. Lim, W.S., et al., BTS guidelines for the management of community acquired pneumonia in adults: update 2009. Thorax, 2009. 64(Suppl 3): p. iii1–iii55.

3. Torres, A., et al., International ERS/ESICM/ESCMID/ALAT guidelines for the management of hospital-acquired pneumonia and ventilator-associated pneumonia: Guidelines for the management of hospital-acquired pneumonia (HAP)/ventilator-associated pneumonia (VAP) of the European Respiratory Society (ERS), European Society of Intensive Care Medicine (ESICM), European Society of Clinical Microbiology and Infectious Diseases (ESCMID) and Asociación Latinoamericana del Tórax (ALAT). Eur Respir J, 2017. 50(3).

4. Luna, C.M., et al., Is a strategy based on routine endotracheal cultures the best way to prescribe antibiotics in ventilator-associated pneumonia? Chest, 2013. 144(1): p. 63–71.

5. Musher, D.M., et al., Can an etiologic agent be identified in adults who are hospitalized for community-acquired pneumonia: results of a one-year study. J Infect, 2013. 67(1): p. 11–8.

6. Akram, A.R., et al., In situ identification of Gram-negative bacteria in human lungs using a topical fluorescent peptide targeting lipid A. Sci Transl Med, 2018. 10(464).

7. Curran, J., et al., Estimating daily antibiotic harms: an umbrella review with individual study meta-analysis. Clin Microbiol Infect, 2022. 28(4): p. 479–490.

8. Tian, S., et al., The role of confocal laser endomicroscopy in pulmonary medicine. Eur Respir Rev, 2023. 32(167).

9. Mills, B., et al., Molecular detection of Gram-positive bacteria in the human lung through an optical fiber-based endoscope. Eur J Nucl Med Mol Imaging, 2021. 48(3): p. 800–807.

10. Craven, T.H., et al., Activated neutrophil fluorescent imaging technique for human lungs. Sci Rep, 2021. 11(1): p. 976.

11. Torrado, B., et al., Fluorescence lifetime imaging microscopy. Nature Reviews Methods Primers, 2024. 4(1).

12. Williams, E., et al., High speed spectral fluorescence lifetime imaging for life science applications. SPIE BiOS. Vol. 10889. 2019: SPIE.

13. Humphries, D.C., et al., Specific in situ immuno-imaging of pulmonary-resident memory lymphocytes in human lungs. Frontiers in Immunology, 2023. 14.

14. Stone, J.M., et al., Low index contrast imaging fibers. Optics Letters, 2017. 42(8): p. 1484–1487.

15. Kuhns, D.B., et al., Isolation and Functional Analysis of Human Neutrophils. Curr Protoc Immunol, 2015. 111: p. 7.23.1–7.23.16.

16. Bankhead, P., et al., QuPath: Open source software for digital pathology image analysis. Sci Rep, 2017. 7(1): p. 16878.

17. Conway Morris, A., et al., 16S pan-bacterial PCR can accurately identify patients with ventilator-associated pneumonia. Thorax, 2017. 72(11): p. 1046–1048.

18. Lakowicz, J., et al., FLUORESCENCE LIFETIME IMAGING. Analytical Biochemistry, 1992. 202(2): p. 316–330.

19. Van Munster, E. and T. Gadella, φFLIM:: a new method to avoid aliasing in frequency-domain fluorescence lifetime imaging microscopy. Journal of Microscopy-Oxford, 2004. 213: p. 29–38.

20. Craig, A., et al., Neutrophil recruitment to the lungs during bacterial pneumonia. Infect Immun, 2009. 77(2): p. 568–75.

21. Perperidis, A., et al., Automated Detection of Uninformative Frames in Pulmonary Optical Endomicroscopy. IEEE Trans Biomed Eng, 2017. 64(1): p. 87–98.

22. Gouzou, D., et al., Applications of machine learning in time-domain fluorescence lifetime imaging: a review. Methods Appl Fluoresc, 2024. 12(2).

23. Haloubi, T., et al., Motion Compensation in Pulmonary Fluorescence Lifetime Imaging: An Image Processing Pipeline for Artefact Reduction and Clinical Precision. Open Journal of Engineering in Medicine and Biology, 2025: p. 1–11.

24. Walsh, A.J., et al., Temporal binning of time-correlated single photon counting data improves exponential decay fits and imaging speed. Biomed Opt Express, 2016. 7(4): p. 1385–99.

