## Supplementary material for "Endomicroscopic fluorescence lifetime imaging enables molecular detection and targeted sampling in the distal human lung": FigS1_Static with description

### Fig S1

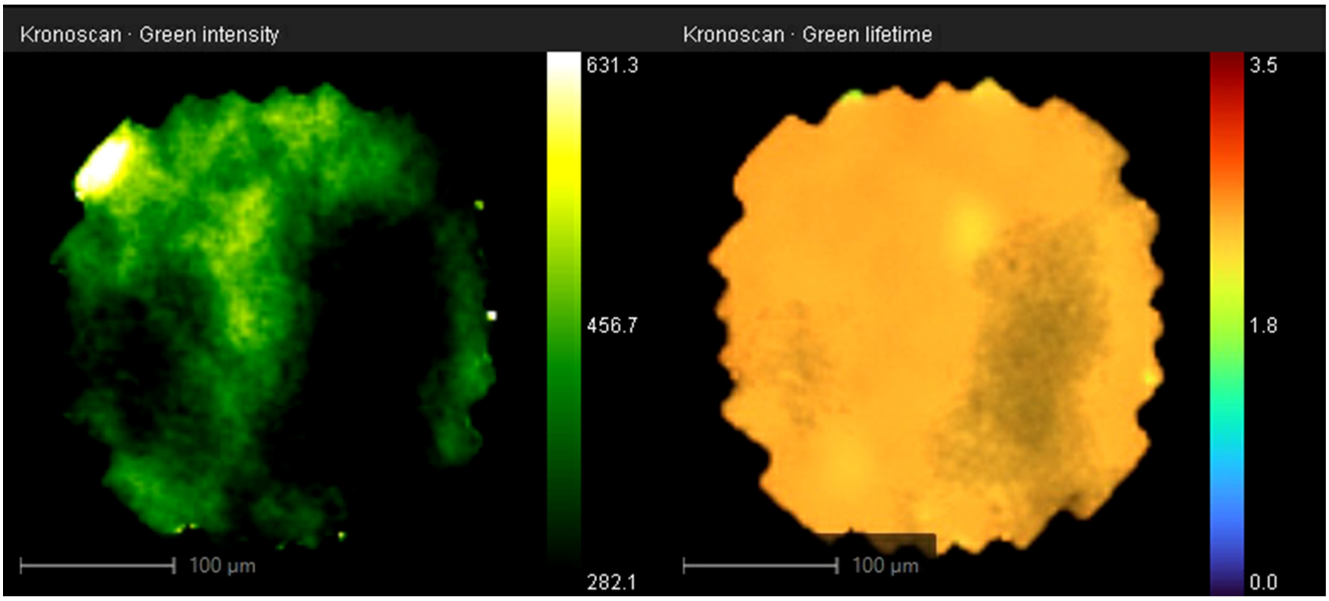


Fig S1

**Normal lung tissue visualised using eFLIM system.** Video was obtained with an imaging fibre deployed through the working channel of a bronchoscope in ex-vivo human lungs, illustrating the optical fibre entering the alveolar space (around 00:20.38). All images were obtained with spectral emission range 498 – 565 nm. The left-hand image shows fluorescence intensity (autoscaled) and the right-hand image shows fluorescence lifetime (scale 1 – 3.5 ns). Representative video from a single lung.
