## Supplementary material for "Endomicroscopic fluorescence lifetime imaging enables molecular detection and targeted sampling in the distal human lung": FigS2_Static with description

### Fig S2

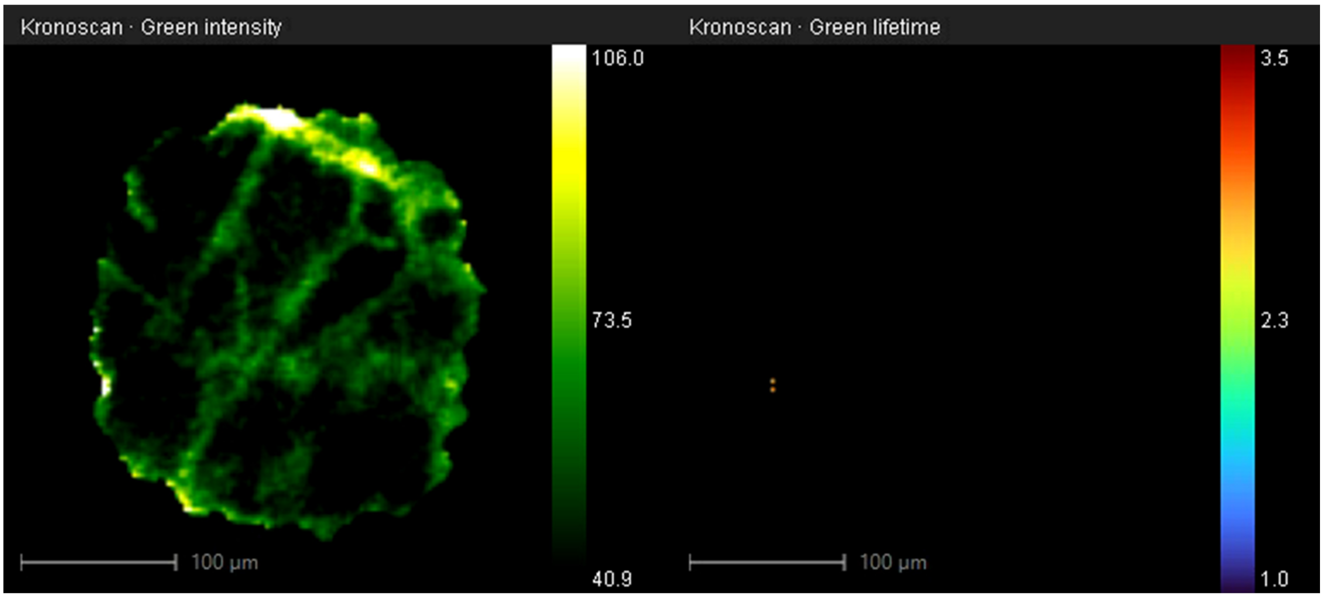


Fig S2

**Figure S2. SmartProbe delivery into the alveolar space visualised using eFLIM.** Video obtained with an imaging fibre deployed through the working channel of a bronchoscope in ex-vivo human lungs following confirmation of alveolar access by characteristic alveolar microanatomy on eFLIM imaging. Instillation of 100 µL SmartProbe into the alveolar space is indicated by transient image distortion and the appearance of fluid microbubbles within the imaging field. All images were obtained with spectral emission range 498–565 nm. The left-hand image shows fluorescence intensity (autoscaled) and the right-hand image shows fluorescence lifetime (scale 1–3.5 ns). Representative video from a single lung.
