## Supplementary figures and images for "Endomicroscopic fluorescence lifetime imaging enables molecular detection and targeted sampling in the distal human lung"

### FigS3

#### Fig S3

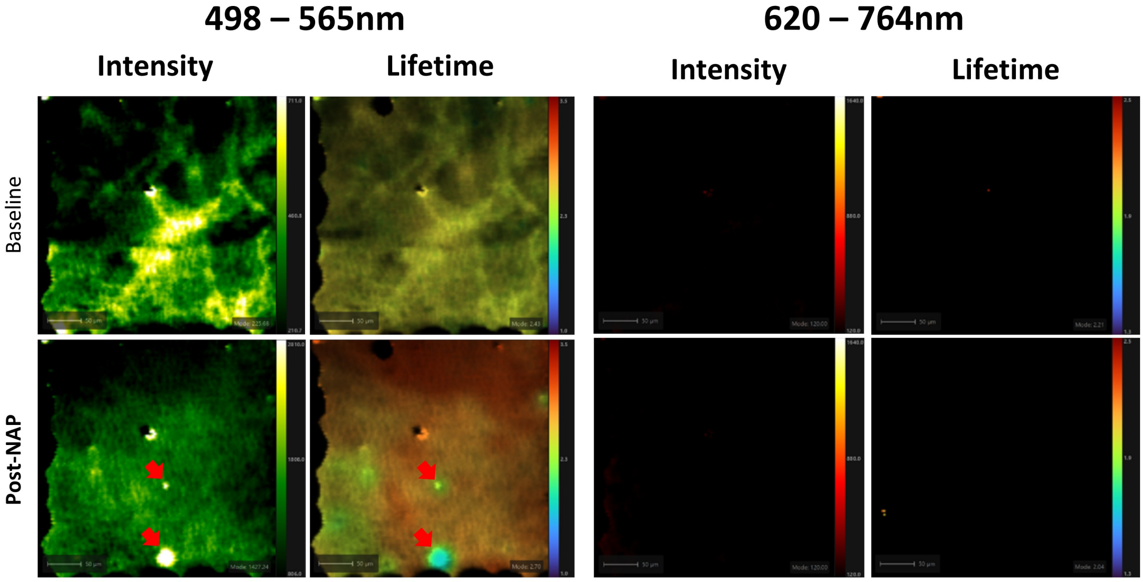


Fig S3. Representative ex-vivo images with NAP including Band Two wavelength.

### FigS4

#### Fig S4

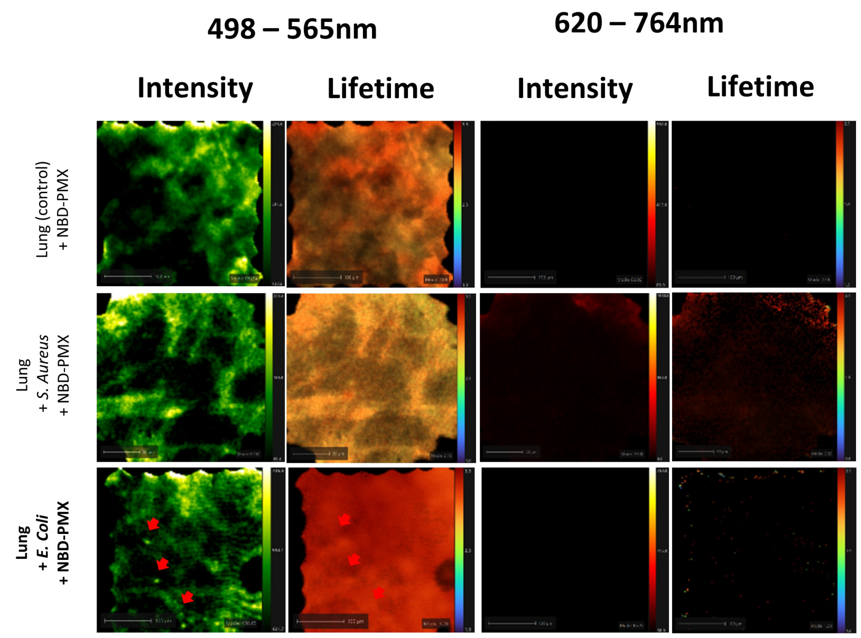


Fig S4. Representative ex-vivo images with NBD-PMX including Band Two wavelength.
