## Supplementary material for "Endomicroscopic fluorescence lifetime imaging enables molecular detection and targeted sampling in the distal human lung": FigS5_Static with description

### Figure S5

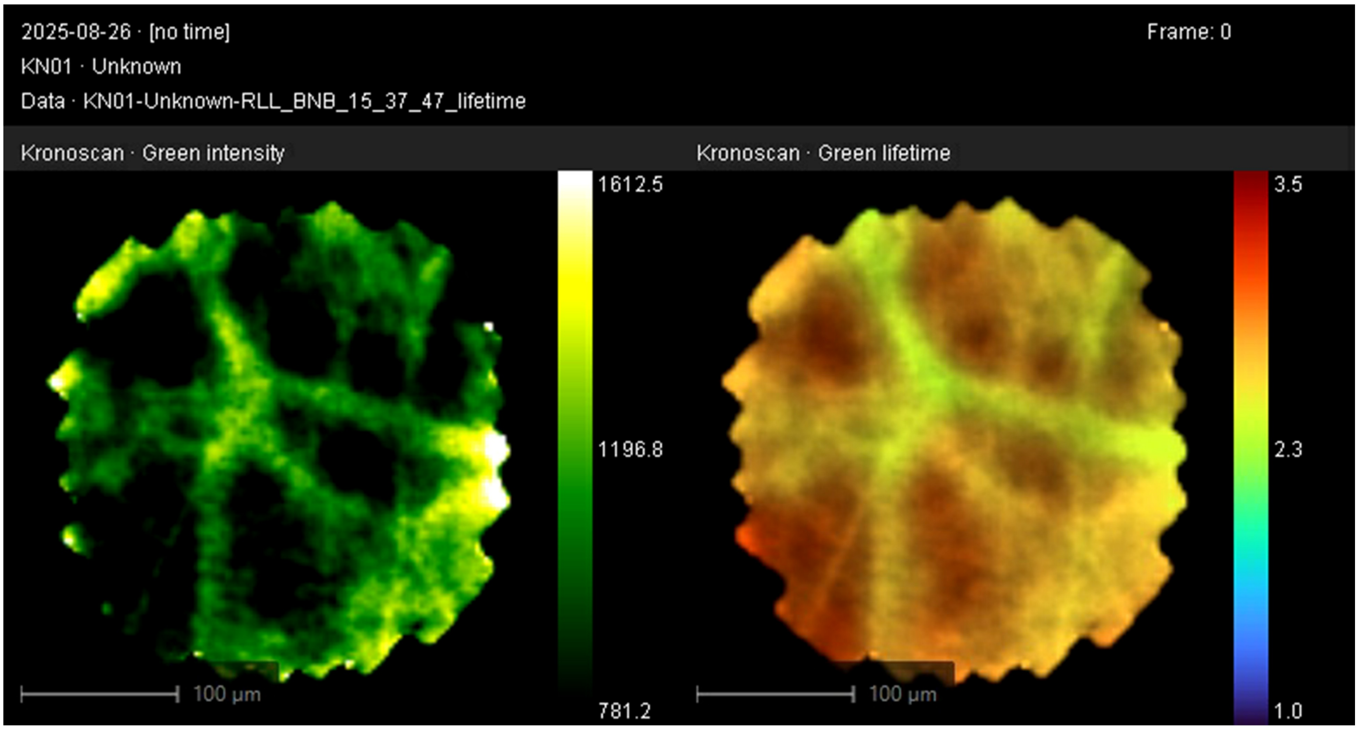


Fig S5. **Representative ex-vivo video after instillation of activated neutrophils followed by instillation of NAP**

Video was obtained with an imaging fibre deployed through the working channel of a bronchoscope in ex-vivo human lungs. Characteristic NAP signal observed from approximately 0:20. All images were obtained with spectral emission range 498 – 565 nm. The left-hand image shows fluorescence intensity (autoscaled) and the right-hand image shows fluorescence lifetime (scale 1 – 3.5 ns). Representative video from a single lung.
