## Supplementary Table T1 for "Endomicroscopic fluorescence lifetime imaging enables molecular detection and targeted sampling in the distal human lung"

| Test | Panoptes performance | | EoT performance | | Notes |
|---|---|---|---|---|---|
| **1. Ability to pass through working channel of bronchoscope while maintaining imaging functionality.** | - Regular Ambu aScope 4<br>- Large Ambu aScope 4<br>- Olympus mobilescope | P<br>P<br>P | - Regular Ambu aScope 4<br>- Large Ambu aScope 4<br>- Olympus mobilescope | F<br>P<br>P | EoT appears to stick in regular Ambu aScope 4. |
| **2. Sufficiently rigid to achieve transbronchial pass into alveolar space.** | Pass | | Pass | | Subjectively more tactile feedback when passing into alveolar space with EoT. |
| **3. Lung damage observed following transbronchial passage compared to commercially available device.** | Pass | | Pass | | Acceptable damage comparable to commercially available device, Alveoflex. |
| **4. Achieve images of alveolar space.** | Pass | | Pass | | Operators felt Panoptes images were potentially sharper. |
| **5. Achieve images of pleura.** | Pass | | Pass | | |
| **6. Achieve images of bronchi.** | Pass | | Pass | | |
| **7. Ability to access 3 subsections of right upper lobe.** | Fail | | Pass | | Difficulty in advancing Panoptes into > 2 subsections of upper lobe, resulting in fibre breakage. |
| **8. Ability to access 3 subsections of right middle lobe.** | Pass | | Pass | | |
| **9. Ability to access 3 subsections of right lower lobe.** | Pass | | Pass | | |
| **10. Ability to access 3 subsections of left upper lobe.** | Fail | | Pass | | Difficult in advancing Panoptes into > 2 subsections as above. |
| **11. Ability to access 3 subsections of left lower lobe.** | Pass | | Pass | | |
| **12. Bend radius within bronchoscope.** | Limited to 120° | | Able to achieve maximal bend radius of bronchoscope (180°) | | |
| **13. Remain usable for duration of imaging session (approx. 20 minutes)** | Pass | | Pass | | |
| **14. Ability to delivery SmartProbes into lung.** | Pass | | Pass | | EoT requires larger air flush of 2ml due to larger capillary size. |
| **15. Ability to aspirate fluid via working channel.** | Variable success – achieving aspirates of 0 – 100 μL | | Pass. Achieved aspirates of 115 – 586 μL. | | |

Table T1:

Comparison of performance of the imaging fibres Eyes on Target (EoT) and previous fibre model (Panoptes) [9]
